# Gene model for the ortholog of *slmb* in *Drosophila pseudoobscura*

**DOI:** 10.64898/2026.01.07.698268

**Authors:** Megan E. Lawson, Aidan Long, Chase Poulsen, Joseph Zajac, Lindsey J. Long, Chinmay Rele, Christine M. Fleet

## Abstract

Gene model for the ortholog of supernumerary limbs (*slmb*) in the Apr. 2013 (BCM-HGSC Dpse_3.0/DpseGB3) Genome Assembly (GenBank Accession: GCA_000001765.2) of *Drosophila pseudoobscura*. This ortholog was characterized as part of a developing dataset to study the evolution of the Insulin/insulin-like growth factor signaling pathway (IIS) across the genus *Drosophila* using the Genomics Education Partnership gene annotation protocol for Course-based Undergraduate Research Experiences.

## Introduction

*This article reports a predicted gene model generated by undergraduate work using a structured gene model annotation protocol defined by the Genomics Education Partnership (GEP; thegep.org) for Course-based Undergraduate Research Experience (CURE). The following information in quotes may be repeated in other articles submitted by participants using the same GEP CURE protocol for annotating Drosophila species orthologs of* Drosophila melanogaster *genes in the insulin signaling pathway*.

“In this GEP CURE protocol students use web-based tools to manually annotate genes in non-model Drosophila species based on orthology to genes in the well-annotated model organism fruitfly *Drosophila melanogaster*. The GEP uses web-based tools to allow undergraduates to participate in course-based research by generating manual annotations of genes in non-model species (Rele et al., 2023). Computational-based gene predictions in any organism are often improved by careful manual annotation and curation, allowing for more accurate analyses of gene and genome evolution (Mudge and Harrow 2016; Tello-Ruiz et al., 2019). These models of orthologous genes across species, such as the one presented here, then provide a reliable basis for further evolutionary genomic analyses when made available to the scientific community.” (Myers et al., 2024).

“The particular gene ortholog described here was characterized as part of a developing dataset to study the evolution of the Insulin/insulin-like growth factor signaling pathway (IIS) across the genus Drosophila. The Insulin/insulin-like growth factor signaling pathway (IIS) is a highly conserved signaling pathway in animals and is central to mediating organismal responses to nutrients (Hietakangas and Cohen 2009; Grewal 2009).” (Myers et al., 2024).

The gene *slmb* (*slimb, ica, shiva, MENE(3R)-B*, or *supernumerary limbs*) was identified in *D. melanogaster* by a genetic screen for genetic mutations affecting body patterning (Jiang and Struhl 1998). The gene product is an F-box/WD40-repeat protein that acts as a substrate adaptor of a conserved SCF (Skp1-Cullin-F-box) E3 ubiquitin ligase, targeting its interacting partners for proteasomal degradation (Jiang and Struhl 1998; Bocca et al., 2001). Reported mutants are hypomorphic or lethal, with ectopic activation of the Hedgehog (Hh) signaling pathway. Thus, wild-type slmb is proposed to act as a negative regulator of this pathway in the absence of ligand (Jiang and Struhl 1998). During the maturation of the nervous system, slmb interacts with Akt, an activator of the insulin signaling pathway, and promotes Akt ubiquitination, thereby inactivating the Insulin Receptor/Phosphoinositide 3-Kinase/Target Of Rapamycin pathway and facilitating dendrite pruning in the ddaC neurons of the peripheral nervous system (Wong et al., 2013). slmb is also known to function in the establishment of axis specification during limb development (Theodosiou et al., 1998), apicobasal polarity of epithelial cells (Skwarek et al., 2014), centrosome development (Wojcik et al., 2000), and copper homeostasis (Zhang et al., 2020).

“*D. pseudoobscura* is part of the *pseudoobscura* species subgroup within the *obscura* species group in the subgenus *Sophophora* of the genus *Drosophila* (Sturtevant 1942; Buzzati-Traverso and Scossiroli 1955). It was first described by Frolowa (1929). The *pseudoobscura* species subgroup is endemic to the western hemisphere, where *D. pseudoobscura* is distributed throughout Western North America, Mexico, and Central America (Markow and O’Grady 2005). An additional population of *D. pseudoobscura*, found near Bogota, Colombia, is partially reproductively isolated from the North and Central American populations (Prakash 1972). *D. pseudoobscura* is found primarily in chaparral and temperate forests. *D. pseudoobscura* has been studied extensively in the context of ecological and behavioral genetics, speciation, and genome evolution (Powell 1997).” (Lawson et al.).

We propose a gene model for the *D. pseudoobscura* ortholog of the *D. melanogaster supernumerary limbs* (*slmb*) gene. The genomic region of the ortholog corresponds to the uncharacterized protein LOC6897521 (RefSeq accession XP_002137670.3) in the Apr. 2013 (BCM-HGSC Dpse_3.0/DpseGB3) Genome Assembly of *D. pseudoobscura* (GenBank Accession: GCA_000001765.2). This model is based on RNA-Seq data from *D. pseudoobscura* (SRP006203-Chen et al., 2014) and *slmb* in *D. melanogaster* using FlyBase release FB2023_02 (GCA_000001215.4; Larkin et al., 2021; Gramates et al., 2022; Jenkins et al., 2022).

### Synteny

The target gene, *slmb*, occurs on chromosome 3R in *D. melanogaster* and is flanked upstream by *CG5793* and *Odorant-binding protein 93a* (*Obp93a*) and downstream by *Bride of doubletime* (*Bdbt*) and peter pan (*ppan*). The *tblastn* search of *D. melanogaster* slmb-PA (query) against the *D. pseudoobscura* (GenBank Accession: GCA_000001765.2) Genome Assembly (database) placed the putative ortholog of *slmb* within scaffold CM000070 (CM000070.3) at locus LOC6897521 (XP_002137670.3)— with an E-value of 0.0 and a percent identity of 95.65%. Furthermore, the putative ortholog is flanked upstream by LOC6897520 (XP_033241694.1) and LOC4802242 (XP_001359203.2), which correspond to *CG5793* and *Obp93a* in *D. melanogaster* (E-value: 7e-135 and 3e-101; identity: 81.82% and 69.04%, respectively, as determined by *blastp*; Figure 1A, Altschul et al., 1990). The putative ortholog of *slmb* is flanked downstream by LOC4802245 (XP_015037694.2) and LOC4802246 (XP_001359205.2), which correspond to *Bdbt* and *ppan* in *D. melanogaster* (E-value: 3e-171and 0.0; identity: 76.92% and 81.64%, respectively, as determined by *blastp*). The putative ortholog assignment for *slmb* in *D. pseudoobscura* is supported by the following evidence: synteny in the genomic neighborhoods of both species is completely conserved, and all *BLAST* results are of very high quality for both the target gene and neighboring genes.

**Figure 1.**
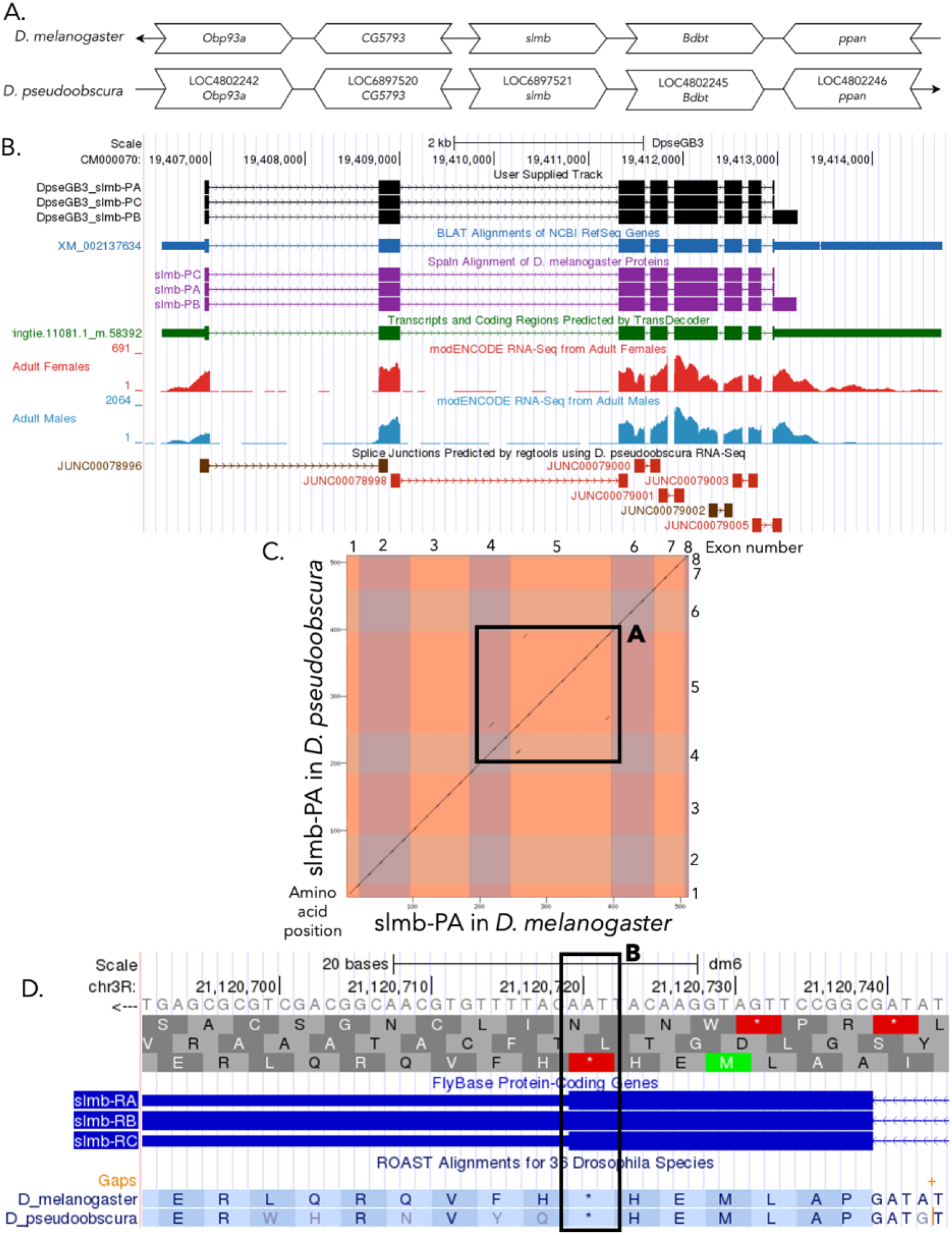
*slmb* gene model comparison between *Drosophila pseudoobscura* and *Drosophila melanogaster* orthologs. **(A) Synteny comparison of the genomic neighborhoods for *slmb* in *Drosophila melanogaster* and *D. pseudoobscura*.** Thin underlying arrows indicate the DNA strand within which the target gene–*slmb*–is located in *D. melanogaster* (top) and *D. pseudoobscura* (bottom). Thin arrow pointing to the right indicates that *slmb* is on the positive (+) strand in *D. pseudoobscura*, and thin arrow pointing to the left indicates that *slmb* is on the negative (-) strand in *D. melanogaster*. The wide gene arrows pointing in the same direction as *slmb* are on the same strand relative to the thin underlying arrows, while wide gene arrows pointing in the opposite direction of *slmb* are on the opposite strand relative to the thin underlying arrows. White gene arrows in *slmb* indicate orthology to the corresponding gene in *D. melanogaster*. Gene symbols given in the *D. pseudoobscura* gene arrows indicate the orthologous gene in *D. melanogaster*, while the locus identifiers are specific to *D. pseudoobscura*. **(B) Gene Model in GEP UCSC Track Data Hub** (Raney et al., 2014). The coding-regions of *slmb* in *D. pseudoobscura* are displayed in the User Supplied Track (black); coding exons are depicted by thick rectangles and introns by thin lines with arrows indicating the direction of transcription. Subsequent evidence tracks include BLAT Alignments of NCBI RefSeq Genes (dark blue, alignment of Ref-Seq genes for *D. pseudoobscura*), Spaln of D. melanogaster Proteins (purple, alignment of Ref-Seq proteins from *D. melanogaster*), Transcripts and Coding Regions Predicted by TransDecoder (dark green), RNA-Seq from Adult Females and Adult Males (red and light blue, respectively; alignment of Illumina RNA-Seq reads from *D. pseudoobscura*), and Splice Junctions Predicted by regtools using *D. pseudoobscura* RNA-Seq (SRP006203). Splice junctions shown have a minimum read-depth of 10 with 500-999 and >1000 supporting reads in brown and red, respectively. **(C) Dot Plot of slmb-PA in *D. melanogaster* (*x*-axis) vs. the orthologous peptide in *D. pseudoobscura* (*y*-axis)**. Amino acid number is indicated along the left and bottom; coding-exon number is indicated along the top and right, and exons are also highlighted with alternating colors. Box A highlights some minor repeats present in the fourth and fifth exons of this gene. **(D) ROAST Alignments for the final exon of *slmb* in *D. pseudoobscura* and *D. melanogaster* shown in UCSC Genome Browser on D. melanogaster Aug. 2014 (BDGP Release 6+ ISO1 MT/dm6)**. The protein alignment track at the bottom shows the aligned amino acid sequence for the shown portion of the final exons of each isoform of *slmb* in *D. melanogaster* and *D. pseudoobscura*. Box B highlights the stop codon that undergoes stop codon readthrough in *D. melanogaster* and is conserved in the *D. pseudoobscura* protein sequence as well.

### Protein Model

*slmb* in *D. pseudoobscura* has three protein-coding isoforms: slmb-PA, slmb-PB, and slmb-PC (Figure 1B). mRNA isoforms slmb-RA and slmb-RC are identical and contain eight protein-coding exons. slmb-RB contains eight protein-coding exons as well, with its only differentiating factor being that it has a longer final exon compared to the other two isoforms. Relative to the ortholog in *D. melanogaster*, the RNA coding-exon number is conserved between these three isoforms in both species. Additionally, in *D. melanogaster*, the longer final exon sequence for slmb-RB is attributed to a stop codon readthrough in Flybase Release FB2023_02, and this may also be true for the *D. pseudoobscura* ortholog of this gene (see “special characteristics of the protein model”; Larkin et al., 2021). The sequence of slmb-PA in *D. pseudoobscura* has 98.82% identity (E-value: 0.0) with the protein-coding isoform slmb-PA in *D. melanogaster*, as determined by *blastp* (Figure 1C). A few minor repeats are present in the fourth and fifth exons of this isoform, but overall this protein alignment is very well-conserved (Figure 1C, Box A). Coordinates of these curated gene models (slmb_PA, slmb_PB, and slmb_PC) are stored by NCBI at GenBank/BankIt (accession BK064647 - BK064648 representing the unique isoforms). This gene model can also be seen within the target genome at this TrackHub.

### Special characteristics of the protein model

#### Stop codon readthrough in the final exon of slmb-RB

slmb-RB in *D. melanogaster* is differentiated from the other isoforms present by its last exon (flybase ID: 9_13730_2), which is longer than the eighth exon found in isoforms slmb-RA and slmb-RC (flybase ID: 8_13730_2). In Flybase Release FB2023_02, this is attributed to a stop codon readthrough characteristic of this protein isoform in *D. melanogaster* (Larkin et al., 2021). Upon analysis of the ROAST Alignments for *D. pseudoobscura* in the *D. melanogaster* genome browser, it appears that this stop codon is conserved in both species, and thus the stop codon readthrough may also occur in the translation of isoform slmb-RB in *D. pseudoobscura* (Figure 1D, Box B).

## Methods

“Detailed methods including algorithms, database versions, and citations for the complete annotation process can be found in Rele et al. (2023). Briefly, students use the GEP instance of the UCSC Genome Browser v.435 (https://gander.wustl.edu; Kent WJ et al., 2002; Navarro Gonzalez et al., 2021) to examine the genomic neighborhood of their reference IIS gene in the *D. melanogaster* genome assembly (Aug. 2014; BDGP Release 6 + ISO1 MT/dm6). Students then retrieve the protein sequence for the *D. melanogaster* reference gene for a given isoform and run it using *tblastn* against their target *Drosophila* species genome assembly on the NCBI BLAST server (https://blast.ncbi.nlm.nih.gov/Blast.cgi; Altschul et al., 1990) to identify potential orthologs. To validate the potential ortholog, students compare the local genomic neighborhood of their potential ortholog with the genomic neighborhood of their reference gene in *D. melanogaster*. This local synteny analysis includes at minimum the two upstream and downstream genes relative to their putative ortholog. They also explore other sets of genomic evidence using multiple alignment tracks in the Genome Browser, including BLAT alignments of RefSeq Genes, Spaln alignment of *D. melanogaster* proteins, multiple gene prediction tracks (e.g., GeMoMa, Geneid, Augustus), and modENCODE RNA-Seq from the target species. Detailed explanation of how these lines of genomic evidence are leveraged by students in gene model development are described in Rele et al. (2023). Genomic structure information (e.g., CDSs, intron-exon number and boundaries, number of isoforms) for the *D. melanogaster* reference gene is retrieved through the Gene Record Finder (https://gander.wustl.edu/∼wilson/dmelgenerecord/index.html; Rele et al., 2023). Approximate splice sites within the target gene are determined using *tblastn* using the CDSs from the *D. melanogaste*r reference gene. Coordinates of CDSs are then refined by examining aligned modENCODE RNA-Seq data, and by applying paradigms of molecular biology such as identifying canonical splice site sequences and ensuring the maintenance of an open reading frame across hypothesized splice sites. Students then confirm the biological validity of their target gene model using the Gene Model Checker (https://gander.wustl.edu/∼wilson/dmelgenerecord/index.html; Rele et al., 2023), which compares the structure and translated sequence from their hypothesized target gene model against the *D. melanogaster* reference gene model. At least two independent models for a gene are generated by students under mentorship of their faculty course instructors. Those models are then reconciled by a third independent researcher mentored by the project leaders to produce the final model. Note: comparison of 5’ and 3’ UTR sequence information is not included in this GEP CURE protocol.” (Gruys et al., 2025)

## Supporting information

Gene model data files

## Supplemental Files

1. Zip file containing a FASTA, PEP, GFF files for the gene model
2. Figure 1 in high resolution

## Metadata

Bioinformatics, Genomics, *Drosophila*, Genotype Data, New Finding

## Acknowledgements

We would like to thank Wilson Leung for developing and maintaining the technological infrastructure that was used to create this gene model and Laura K. Reed for overseeing the project. Thank you to FlyBase for providing the definitive database for *Drosophila melanogaster* gene models. Further, we would like to thank the editors and developers at the journal *microPublication: Biology* for assistance in developing the template for these single gene ortholog publications.

## Funding

This material is based upon work supported by the National Science Foundation (1915544) and the National Institute of General Medical Sciences of the National Institutes of Health (R25GM130517) to the Genomics Education Partnership (GEP; https://thegep.org/; PI-LKR). Any opinions, findings, and conclusions or recommendations expressed in this material are solely those of the author(s) and do not necessarily reflect the official views of the National Science Foundation nor the National Institutes of Health.

## Notes

https://gander.wustl.edu/cgi-bin/hgTracks?db=DpseGB3&lastVirtModeType=default&lastVirtModeExtraState=&virtModeType=default&virtMode=0&nonVirtPosition=&position=CM000070%3A19406434%2D19413707&hgsid=68144_TXlZmmyan8VdhJO71P8VQIR3l6l0

### Competing Interest Statement

The authors have declared no competing interest.

